# Classification of multigene families of African swine fever viruses

**DOI:** 10.1101/2020.02.20.957290

**Authors:** Zhaozhong Zhu, Huiting Chen, Yang Cao, Taijiao Jiang, Yuanqiang Zou, Yousong Peng

**Affiliations:** College of Biology, Hunan University, Changsha, China; Center of Growth, Metabolism and Aging, Key Laboratory of Bio-Resource and Eco-Environment of Ministry of Education, College of Life Sciences, Sichuan University, Chengdu 610065, Sichuan, China; Center for Systems Medicine, Institute of Basic Medical Sciences, Chinese Academy of Medical Sciences & Peking Union Medical College, Beijing 100005; Suzhou Institute of Systems Medicine, Suzhou, Jiangsu 215123, China; Changsha Qiangze Biotech Co., Ltd, Changsha, China

## Abstract

African swine fever virus (ASFV) is a large and complex double-stranded DNA virus that poses serious threats to the pig industry. It is well-accepted that the multigene family (MGF) proteins are extensively distributed in ASFVs and are generally classified into five families, including MGF-100, MGF-110, MGF-300, MGF-360 and MGF-505. Most MGF proteins, however, have not been well characterized and classified within each family. To bridge this gap, this study first classified the MGF proteins into 35 groups based on protein sequence homology. A web server for classifying the MGF proteins was then established and available for free at http://www.computationalbiology.cn/MGF/home.html. Results showed that the genetic diversity of the MGF groups varied widely, mainly due to the occurrence of indels. In addition, the MGF proteins were predicted to have large structural and functional diversity, and the MGF proteins of the same MGF family tended to have similar structure, location and function. Evolutionary analysis revealed the dynamic changes of the MGF proteins in the ASFV genomes, and more than half of MGF groups were presented in all ASFV genomes, which indicated the important role of MGF proteins in ASFVs. Overall, it is expected that the work would not only provide a detailed classification for MGF proteins, but also facilitate further research on MGF proteins.

## 1 Introduction

African swine fever (ASF) is a viral disease in swine that leads to a high mortality in domestic pigs (Parker, Plowright, & Pierce, 1969). The disease has been endemic in sub-Saharan Africa for many years and has spread to many countries in Eastern Europe and Russia in the past decades (Costard, Mur, Lubroth, Sanchez-Vizcaino, & Pfeiffer, 2013; Galindo & Alonso). In the past two years, ASF has spread to China and caused epidemics in almost all provinces of the country (Ge et al.; Li et al., 2019). Despite the growing threat to the world’s pig industry, effective vaccine and drugs have not yet been developed (Teklue, Sun, Muhammad, Luo, & Qiu, 2019).

African swine fever virus (ASFV), the pathogen of ASF, is a large double-stranded DNA virus with a genome size of 170 kb to 190kb and is the only member of the Asfarviridae family (Arias, Jurado, Gallardo, Fernández-Pinero, & Sánchez-Vizcaíno, 2018; L. K. Dixon, Chapman, Netherton, & Upton, 2013). ASFV encodes more than 150 proteins, including proteins related to viral transcription and replication, structural proteins, viral enzymes, etc (L. K. Dixon et al., 2013). Unfortunately, over half of these proteins have not been well characterized, especially for the multigene family (MGF) proteins (Keßler et al., 2018). In general, the MGF proteins are mainly encoded at both ends of the genome and are the most abundant proteins in ASFVs (Vydelingum, Baylis, Bristow, Smith, & Dixon, 1993; Yozawa, Kutish, Afonso, Lu, & Rock, 1994). Depending on the size of the MGF proteins, they can be divided into five families, including MGF-100, MGF-110, MGF-300, MGF-360 and MGF-505 (Chapman, Tcherepanov, Upton, & Dixon, 2008; Keßler et al., 2018). Previous studies have shown that MGF proteins play important roles in host viral infection, including the transcription and translation, virulence, immune escape, etc. For example, MGF-360 and MGF-505 genes have been shown to attenuate a highly virulent ASFV isolate (Neilan et al., 2002; Zsak et al., 2001). However, most MGF proteins have not been structurally and functionally characterized. Considering the great diversity of MGF proteins in each MGF family, there is also a lack of further classification of the MGF proteins.

To bridge this gap, this study first classified the MGF proteins based on protein sequence homology. After that, the genetic diversity, structure, function and evolution of MGF proteins were thoroughly investigated. We hypothesized that a more detailed classification for MGF proteins could be obtained to facilitate further research on MGF proteins.

## 2 Materials and Methods

### 2.1 MGF grouping

A total of 1552 MGF proteins encoded in 39 ASFV genomes were adapted from Zhu’s study (Zhu et al., 2019). The MGF proteins were then grouped based on sequence homology and network clustering using OrthoFinder (version 2.2.7) (Emms & Kelly, 2015) with default parameters, resulting in a total of 35 protein groups. The protein sequences of each MGF group were further aligned by MAFFT (version 7) (Katoh & Standley, 2013) with default parameters.

### 2.2 Phylogenetic tree inference and visualization

To build the phylogenetic tree of each MGF family, the protein sequences of each MGF group were aligned by MAFFT (version 7) with default parameters. The MEGA (version X) (Kumar, Stecher, Li, Knyaz, & Tamura, 2018) with default parameters was employed to build a maximum-likelihood phylogenetic tree. Bootstrap analysis was then conducted with 100 replicates. Finally, the phylogenetic tree was visualized using Dendroscope (version 2.7.4) (Huson et al., 2007).

### 2.3 Structure and function analysis of MGF groups

The largest MGF protein within each MGF group was selected as queries to predict the structure and function features of MGF groups using various public tools. Specifically, the disordered regions of MGF proteins were predicted by the DISPRED (Peng, Radivojac, Vucetic, Dunker, & Obradovic, 2006) server (available at http://www.dabi.temple.edu/disprot/predictor.php). The secondary structures of MGF proteins were predicted by PHD (Geourjon & Deleage, 1995) (available at https://npsa-prabi.ibcp.fr/cgi-bin/npsa_automat.pl?page=/NPSA/npsa_phd.html).

Additionally, the transmembrane domain of MGF proteins were predicted using the TMHMM (Möller, Croning, & Apweiler, 2001) server (available at http://www.cbs.dtu.dk/services/TMHMM/), while the signal peptide and subcellular localization of MGF proteins were predicted by the SignaIP (Armenteros et al., 2019) server (available at http://www.cbs.dtu.dk/services/SignalP/) and Cell-PLoc 2.0 (Chou & Shen, 2010) (available at http://www.csbio.sjtu.edu.cn/bioinf/Cell-PLoc-2/), respectively. Finally, the post-translational modification of MGF proteins including the N-linked glycosylation, acetylation, and ubiquitination, were predicted by the NetNGlyc (Gupta, Jung, & Brunak, 2004) server (available at http://www.cbs.dtu.dk/services/NetNGlyc/), NetAcet (Kiemer, Bendtsen, & Blom, 2004) server (available at http://www.cbs.dtu.dk/services/NetAcet/) and iUbiq-Lys (Wang, Xiao, & Chou, 2011) server (available at http://www.jci-bioinfo.cn/iUbiq-Lys).

### 2.4 Statistical analysis

All the statistical analyses were conducted in R (version 3.2.5) (R Core Team, 2013). The wilcox rank-sum test was conducted by the function of *wilcox.test()* in R. The correlation coefficient was conducted by the function of *cor.test()* in R.

### 2.5 Data availability

All the protein sequences of MGF groups used in this study are publicly available at http://www.computationalbiology.cn/MGF/home.html.

## Results

### Classification of MGF proteins in ASFV

The number of groups in each MGF family was identified and shown in Figure 1. Specifically, a total of 3, 7, 2, 13 and 10 groups were identified in MGF families of 100, 110, 300, 360 and 505, respectively. The groups were named after a combination of the name of MGF family and a letter that began with “A” and followed alphabet order as the number of proteins in the group decreased. For example, the MGF groups in the family of MGF-110 were named MGF-110-A ∼ MGF-110-G. In addition, most MGF groups contained 20 to 110 proteins except MGF-505J that only contained two MGF proteins (Table S1). The average number of proteins in all MGF groups was 44.

**Figure 1.**
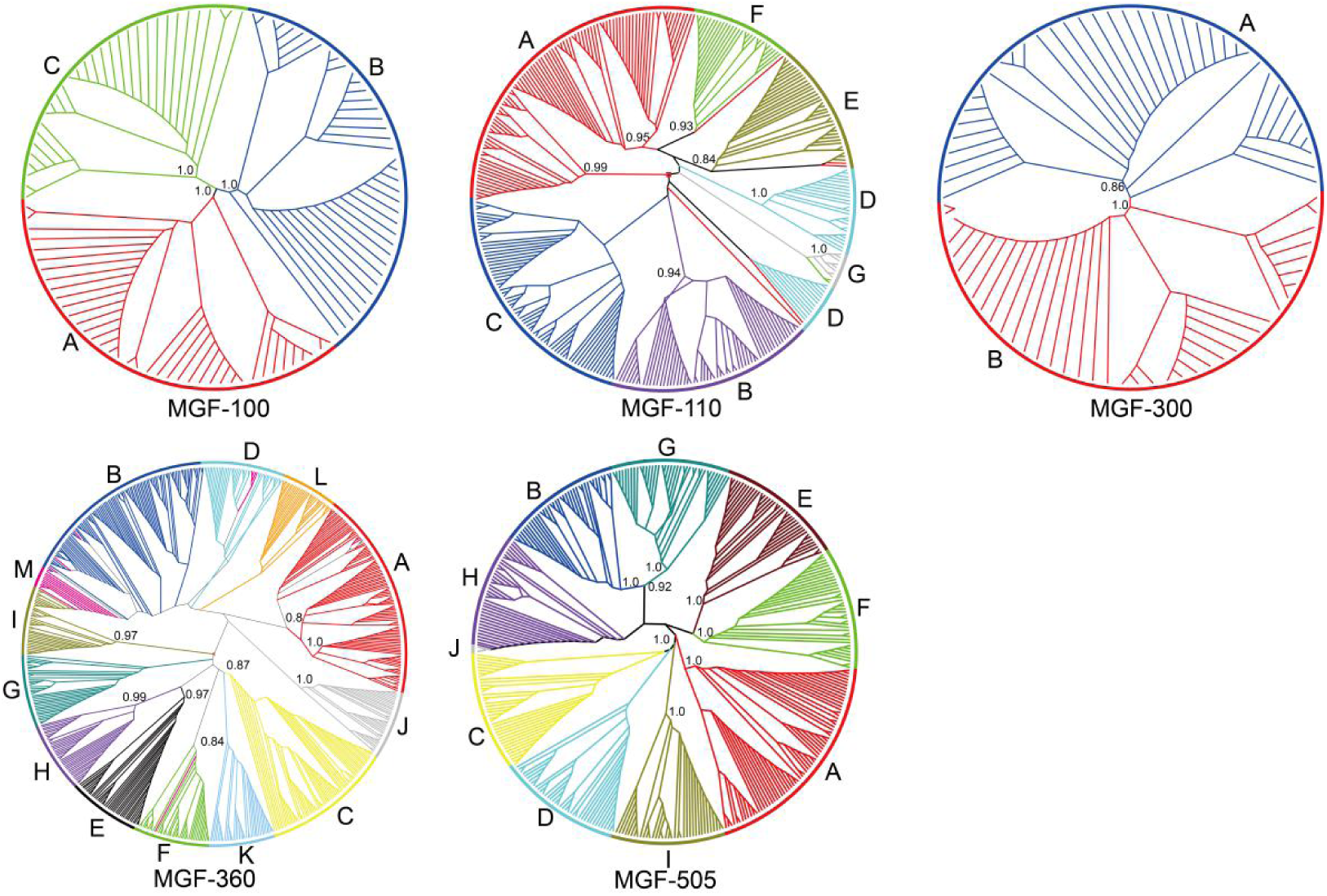
The maximum-likelihood phylogenetic tree for each MGF family. The MGF groups were colored in each MGF family. The letters denoted the names of the MGF groups. The numbers in the tree referred to the bootstrap value.

The maximum-likelihood phylogenetic trees were built for each MGF family based on MGF protein sequences (Figure 1). Results showed that for most MGF groups, the proteins from the same MGF group were clustered together in the phylogenetic trees, indicating the accuracy of the MGF grouping. A leave-one-out test was further performed to validate the MGF grouping. Specifically, each MGF protein was queried against the MGF proteins by blast. The MGF group of the best blast hit (except the query protein) was predicted as the MGF group of the query protein. The predictive accuracy was then calculated for each MGF group. 30 out of 35 groups achieved an accuracy of 100% (Table S2). The remaining MGF groups in MGF-110A, MGF-110E, MGF-360D, MGF-360M and MGF-505J achieved an accuracy of 98%, 97%, 97%, 90% and 50%, respectively.

The 861 MGF proteins downloaded in the NCBI protein database were classified into 34 MGF groups based on the MGF grouping and best blast hit method (Table S3). To facilitate the use of the MGF grouping, a web server named MGFC was established, which is available for free at http://www.computationalbiology.cn/MGF/home.html.

### Genetic diversity of the MGF groups

The genetic diversity of the MGF groups was analyzed. The diversity index, defined as the average ratio of the protein sequence differences, was calculated for each MGF group. Briefly, the value of diversity index ranged from 0.05 to 0.40 with an average of 0.15. While the diversity indexes of some MGF groups were larger than 0.3, such as the MGF-110B and MGF-360B, the diversity indexes of most MGF groups were less than 0.15, especially for the MGF-300 and MGF-505 families (Figure 2).

**Figure 2.**
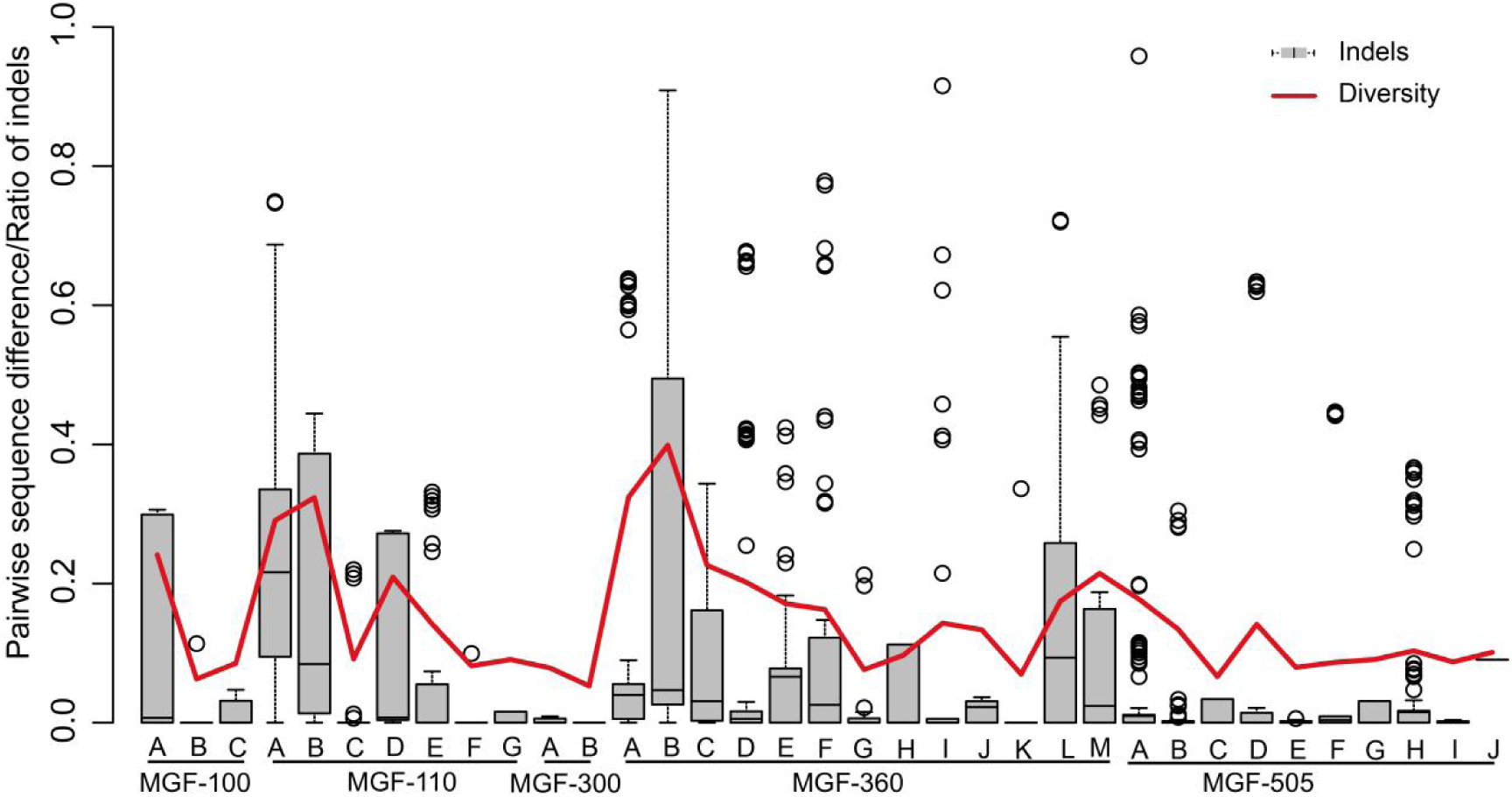
The genetic diversity of the MGF groups. The red curve denoted the diversity index, while the box plot denoted the ratio of indels.

The insertions and deletions (indels) in protein sequences of each MGF group were also investigated. The pairwise ratios of protein sequence difference introduced by indels were calculated for each MGF group. Interestingly, there was a high positive correlation between the diversity index and the average ratio of indels, with a Pearson’s correlation coefficient of 0.82, which indicated that the genetic diversity within the MGF groups was mainly contributed by indels. For example, the largest diversity index of the MGF-360B was 0.40, and the corresponding ratio of indels was also large.

### Structure and function analysis of the MGF groups

We further characterized the structure and function of the MGF proteins for the MGF groups. Nine MGF groups were predicted to be all-alpha proteins, of which eight belonged to MGF-505 family. Only two MGF groups, i.e., the MGF-100C and MGF-300B, were predicted to be alpha-beta proteins. Analysis of structural flexibility showed that eight MGF groups were locally disordered, including all three MGF groups in MGF-100 family.

**Figure 3.**
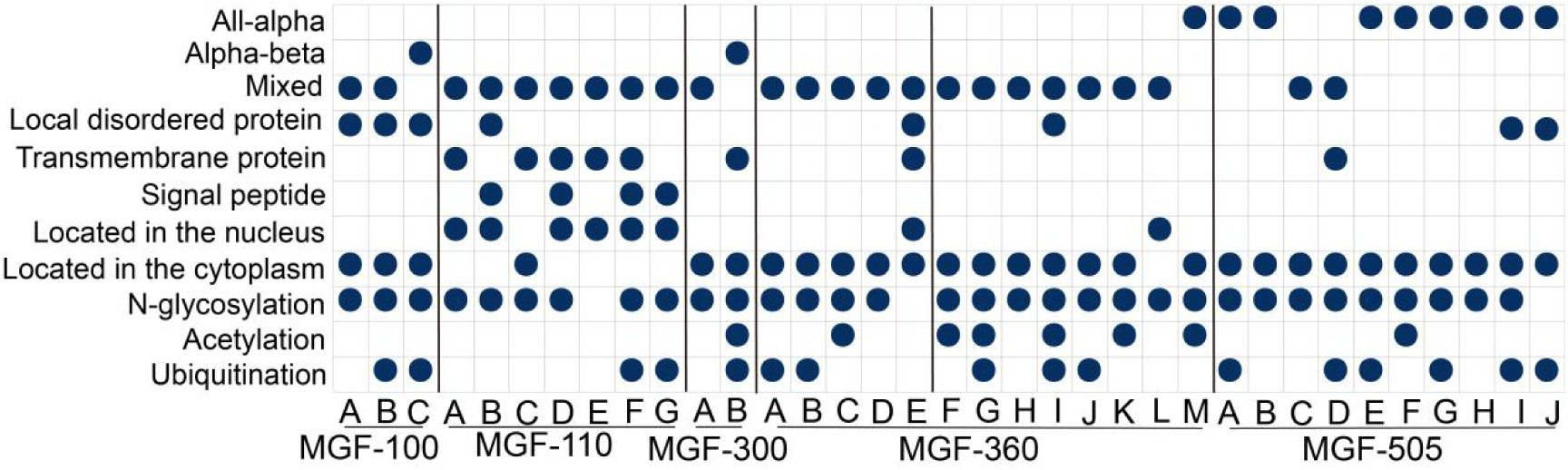
Structure and function analysis of MGF proteins for the MGF groups. In this study, a local disordered protein was defined as a long disordered region containing more than 30 residues (Singh, Ganapathi, & Dash, 2007). In addition, the protein class was determined based on Geourjon’s study (Geourjon & Deleage, 1995). Specifically, a class with an alpha-helix (H) ratio > 45% and a beta-sheet ration (E) < 5% was defined as all-alpha, while a class with H < 5% and E > 45% was defined as all-beta. The alpha-beta class was with H > 30% and E > 20%, and the rest was defined as mixed class.

Eight MGF groups were predicted to contain one or more transmembrane helixes, five of which belonged to the MGF-110 family. Additionally, all the four MGF groups predicted to contain the signal peptide, and six of the eight MGF groups predicted to be located in the nucleus, also belonged to the MGF-110 family, indicating that most MGF groups in the MGF-110 family may be located in both the cell membrane and nucleus. Overall, the majority of the MGF groups were predicted to be located in the cytoplasm.

Post-translational modification of the MGF proteins was analyzed. The N-linked glycosylation was predicted to occur in 32 of the 35 MGF groups. The ubiquitination was also predicted to occur in half of all the MGF groups, while the acetylation was predicted to mainly occur in the MGF groups of the MGF-360 family.

### Evolution of MGF proteins

Finally, we investigated the dynamic evolution of the MGF proteins. The number of the MGF proteins in each MGF group in the ASFVs isolated from 1950 to 2019 was shown in Figure 4. Interestingly, most MGF groups had only one member in an ASFV genome, though some MGF groups had multiple members in an ASFV genome, such as MGF-110A, MGF-360A and MGF-360B. While the number of MGF proteins in an ASFV genome varied, 17 of the 35 MGF groups were present in all ASFVs, indicating the important role of these MGF groups in ASFVs. In addition, some MGF groups presented significant expansion and contraction during the evolution. For example, the MGF-110A had three to five members in the X and IX genotype strains, and no more than two members in eight of the nine viral strains of the genotype I. Moreover, we also found that the viral strain Tengani was lost in the MGF-110A. While some MGF groups were generated in the evolution of ASFVs, such as MGF-360L and MGF-360M, some MGF groups were also lost in the evolution of ASFVs, such as MGF-110G.

**Figure 4.**
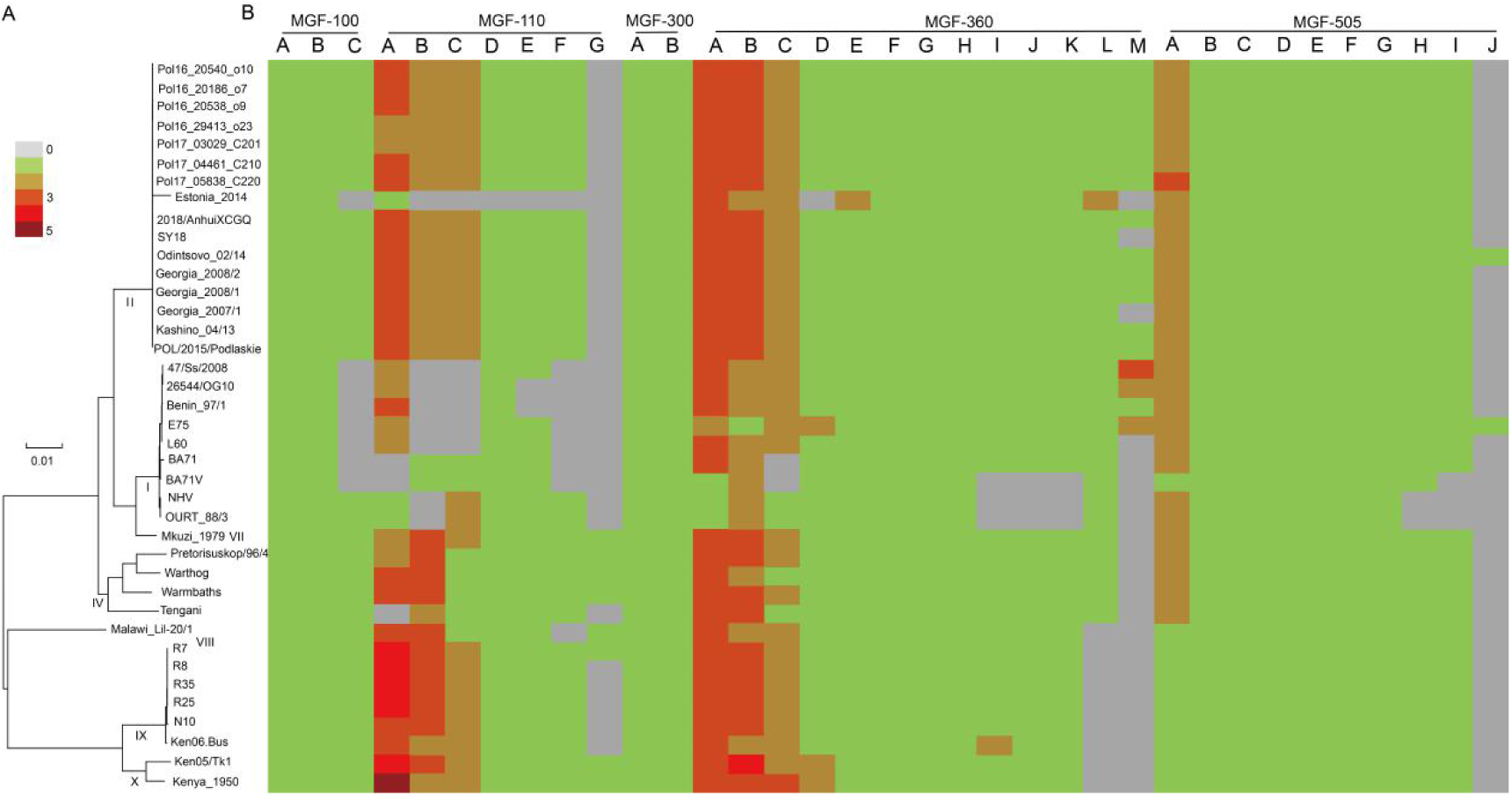
The evolution of the MGF proteins from 1950 to 2019. (A) The maximum-likelihood phylogenetic tree of ASFVs based on the genome sequences adapted from Figure 2A in Zhu’s work (Zhu et al., 2019). The genotypes of the ASFVs were denoted by the bold and italic numbers, while the scale bar represented the number of nucleotide substitutions per site. (B) The number of the MGF proteins in each MGF group in ASFV genomes.

## DISCUSSION

In this study, we first validated the large genetic diversity of the MGF proteins by classifying the MGF proteins of ASFVs into 35 groups. Based on the result of MGF grouping, we systematically analyzed the genetic diversity, structure, function and evolution of the MGF proteins. Interestingly, a strong positive correlation was found between the genetic diversity and the level of indels within the MGF groups, demonstrating that the indels account for the majority of the genetic diversity in MGF groups. This finding is consistent with our previous study that the genetic diversity of ASFV genomes was mainly caused by indels instead of mutations (Zhu et al., 2019).

The MGF groups were predicted to have large structural and functional diversity in our study. This may be related to the diverse functions of MGF proteins reported in previous studies (L. Dixon, Islam, Nash, & Reis, 2019; Netherton et al., 2019). For example, it is reported that the MGF proteins were associated with the virulence, antigenicity and immune escape of ASFVs (Burrage, Lu, Neilan, Rock, & Zsak, 2004; Golding et al., 2016). Our findings suggested that MGF proteins sharing the same MGF family tend to have similar structures, locations and functions. For example, most MGF groups in the MGF-110 family were found to have a mixed composition of secondary structures, transmembrane helixes, signal peptides and N-linked glycosylation, which may imply the same origin of MGF groups within each MGF family.

More than half of MGF groups presented significant expansion and contraction in ASFV genomes. The dynamic changes in the MGF proteins may cause the phenotypic changes in ASFVs, such as the changes in antigen and virulence (Chapman et al., 2008). Many attempts to develop effective vaccines against ASFVs have failed, one possible reason of which is the complex composition of antigens (O’Donnell et al., 2015; Rock, 2017). Therefore, understanding the function and the mechanisms underlying the dynamic changes in MGF proteins may facilitate the development of vaccines and drugs against ASFVs. Taken together, we expect this work would not only provide a classification for MGF proteins, but also facilitate further research on MGF proteins.

## Acknowledgements

This work was supported by the National Key Plan for Scientific Research and Development of China (2016YFD0500300), Hunan Provincial Natural Science Foundation of China (2018JJ3039), the National Natural Science Foundation of China (31671371), the Chinese Academy of Medical Sciences (2016-I2M-1-005), and the Key R&D Program of Sichuan (2018NZ0151).

The authors have declared that no competing interests exist.

## Ethical Statement

Not applicable because no human or animal samples were collected in this study.

